# SMbiot: A Shared Latent Model for Microbiomes and their Hosts

**DOI:** 10.1101/2022.10.28.514090

**Authors:** Madan Krishnamurthy, Lukas Herron, Dwi Susanti, Alyssa Volland-Munson, Germán Plata, Purushottam Dixit

**Affiliations:** Discovery Research, Elanco Animal Health, Greenfield, IN, 46140; Department of Physics, University of Florida, Gainesville, FL 32611; Institute of Physical Sciences and Technology, University of Maryland, College Park, MD 20742; Discovery Research, BiomEdit, LLC., Fishers, IN 46037; Compute & Analytics, BiomEdit, LLC., Fishers, IN 46037; Department of Chemical Engineering, University of Florida, Gainesville, FL 32611; Genetics Institute, University of Florida, Gainesville, FL 32611

## Abstract

The collective nature of the variation in host associated microbial communities suggest that they exhibit low dimensional characteristics. To identify these lower dimensional descriptors, we propose SMbiot (pronounced SIM BY OT): a **S**hared Latent **M**odel for Micro**bio**mes and their hos**t**s. In SMbiot, latent variables embed host-specific microbial communities in a lower dimensional space and the corresponding features reflect controlling axes that dictate community compositions. Using data from different animal hosts, organ sites, and microbial kingdoms of life, we show that SMbiot identifies a small number of host-specific latent variables that accurately capture the compositional variation in host associated microbial communities. By using the same latents to describe hosts’ phenotypic states and the host-associated microbiomes, we show that the latent space embedding is informed by host physiology as well as the associated microbiomes. Importantly, SMbiot enables the quantification of host phenotypic differences associated with altered microbial community compositions in a host-specific manner, underscoring the context specificity of host-microbiome associations. SMbiot can also predict missing host metadata or microbial community compositions. This way, SMbiot is a concise quantitative method to understand the low dimensional collective behavior of host-associated microbiomes.

## Introduction

Animal hosts exist in close association with multiple, species-rich, and functionally versatile microbial communities. These communities span all kingdoms of microbial life including bacteria^1^, archaea^2^, fungi^3^, protozoa^4^, and viruses^5^. These Host-associated communities are recognized to play important roles in human health^6^, as well as feed efficiency and the environmental footprint of livestock species^7^.

Recent developments in multi-omics^8^ portray a comprehensive picture of the collective behavior within the host-microbiome(s) metacommunity^9^. An emergent feature from studies across several hosts is that the hosts’ phenotypic states and the compositions of the several host-associated microbiomes are tightly interdependent^10^. Moreover, the observed covariations are likely to be context-specific and bidirectional. That is, the contribution of specific microbes to specific host phenotypes depends on the presence of other microbes^11^ and environmental and host genetic factors^12^, and both host and the microbiome can affect each other’s states. Therefore, it is reasonable to expect that the observed collective behavior of components of similar host-microbiome ecosystems are low-dimensional, that is, they can be described by a small number of latent variables.

The assembly rules for the host-microbiome ecosystems are complex, as they include several types of interactions such as competition for common resources^13^, predation^14^, cross-feeding^13^, and interactions with the host immune system^15^. Therefore, bottom-up mechanistic models are not equipped to explain the observed covariation among similar ecosystems and the context-dependence of these interactions. Previous quantitative methods based on correlation-based network inference and regression-based methods can identify specific host-microbe associations and predict host phenotypes using microbial abundances and vice versa. These correlation networks hint towards a low-dimensional description of the host-associated microbial ecosystems but do not explicitly obtain it. In contrast, dimensionality reduction methods such as multidimensional scaling (MDS)^16^ can potentially project microbiome compositions or host phenotypic data onto lower dimensional manifolds. However, these methods are descriptive; the lower dimensional embeddings cannot be used to explain microbial compositions and host phenotypic states. Therefore, current computational approaches cannot identify the controlling variables that describe the collective behavior within the host-microbiome metacommunity.

We recently developed a computational method to embed temporal variability in gut microbiomes using latent space descriptors^17^. This allowed us to capture the collective variability in bacterial abundances in the gut microbiome using a small number of latent space dimensions. Importantly, we found that the latent space descriptors correlated strongly with host characteristics such as diet and antibiotic administration status. Here, we make those observations concrete and significantly generalize that work. Specifically, we hypothesize that the collective variations in hosts’ phenotypic states and the compositions of the several associated microbiomes are governed by a small number of controlling variables.

To identify these lower dimensional representations, we present a new framework, SMbiot: A **S**hared Latent **M**odel for micro**bio**mes and their hos**t**s. Briefly, SMbiot uses a shared latent space representation that simultaneously models available microbiome data, for example, microbiota compositions at different host sites, and/or multidimensional data on the hosts’ phenotypic states. Surprisingly to us, using different types of phenotypic data collected on several animal hosts, as well as compositions of microbiomes across several organ sites and kingdoms of life, we find that SMbiot can accurately capture the variation in these metacommunities using a small number of variables. Moreover, SMbiot obtains an approximate latent space representation of the metacommunities when only partial data are available, thereby allowing to predict unobserved microbiome compositions or phenotypic data. As validation, we use SMbiot to predict changes in the gut metabolome of chickens upon antibiotic treatment starting from 16S rRNA sequencing data, showing that accurate predictions using SMbiot enable biological insights while reducing experimental costs. Importantly, we also show that SMbiot quantifies the co-dependence between microbiome and metadata perturbations at any point in the latent space, thereby quantifying the context-specificity of host-microbiome associations. We believe that SMbiot will be a valuable approach in understanding the collective behaviors in animal hosts and their associated microbiomes.

## Results

### SMbiot: A Shared Latent Model for Microbiomes and their Hosts

Here, we briefly describe SMbiot for a specific use case. See **Supplementary Materials** for a detailed and general description. We start with measurements of *P* phenotypic metadata *m_sp_* (*s* ∈ [1, *S*], *p* ∈ [1, *P*]) and abundances of *0* operational taxonomic units (OTUs) *n_so_* (*s* ∈ [1, *S*],*o* ∈ [1,0]) in one body site (gut, rumen, skin, etc.) of *S* hosts (samples). We model the metadata (assumed to be Z-transformed) as a multivariate Gaussian distribution:

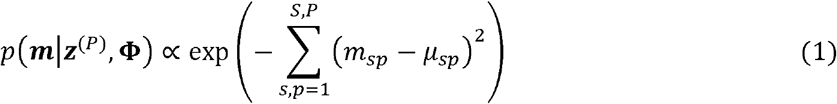

where

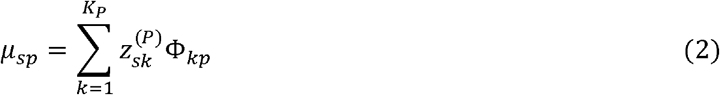

is a low-rank approximation. We choose *K_P_* ≪ *S, P* to impose the low rank. In Eq. (2), ***z***^(*P*)^ are collectively the latents that describe the metadata and **Φ** is the matrix of phenotype-related features. We model microbial abundances using a multinomial distribution,

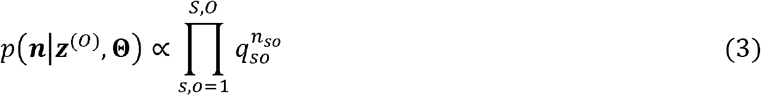

where

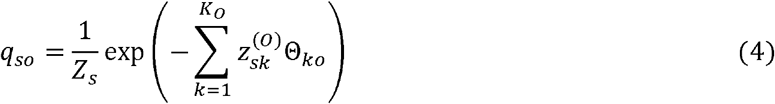

is the reduced-dimensional description of the microbiome^17,18^. In Eq (4) ***z***^(*O*)^ are collectively the latents that describe the microbiome and **Θ** is the matrix of microbiome features. To capture the covariation between host phenotypes and the microbiomes, we use the same *K_c_* ≤ *K_0_, K_P_* latents for the latent space approximation of the microbiome and the phenotypic metadata. We infer the model parameters {***z***^(*P*)^,**z**^(*O*)^**Φ**, **Θ**} by maximizing the combined log-l i ke I hood:

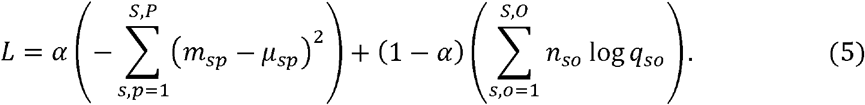

In Eq. (5), 0 < *α* < 1 dictates the relative importance of the two terms.

Other types of information about the host-microbiome metacommunity can be modeled the same way. For example, we may have information about microbiomes at several host sites such as oral swabs, intestine, and feces, in addition to information about the hosts’ phenotypes. In this case, we amend the log-likelihood function in Eq. 5 with several multinomial distributions. If several types of orthogonal metadata are available, for example, fecal and serum metabolomics, we amend the log-likelihood function with additional Gaussian distributions (see **Supplementary Materials**).

Finally, we note that the inferred latent space and the corresponding features (matrices **Θ** and **Φ**) are not uniquely determined but belong to a linear space whose dimension is equal to the dimension of the latent space. Specifically, if we denote by and **Θ**^(**1**)^ and **Θ**^(**2**)^ (and **Φ**^(**1**)^ and **Φ**^(**2**)^) the SMbiot-learnt matrices from two different initializations, there exists a *K × K* invertible matrix ***B*** such that ***Θ***^(**1**)^ = ***B*Θ**^(**2**)^ and **Φ**^(**1**)^ = ***B*Φ**^(2)^. Notably, the inverse of this matrix will also transform the latent space: **Z**^(**1**)^ = **B**^**-1**^Z^(**2**)^.

### SMbiot explains multiple microbiomes associated with the same host using host-specific variables

Microbial species colonize almost every organ of animal hosts^19^. The composition of organ-specific ecosystems can be vastly different, potentially dictated by factors such as nutrient availability and interactions with the host immune system. Yet, given that the ecosystems colonizing the same host share several host-specific characteristics, we tested whether compositional variation in microbiomes colonizing different organ sites of the same host can be explained using host-specific lower dimensional variables.

To that end, we analyzed previously collected data^20^ on chicken bacterial microbiomes sampled from tracheal swabs, the ileum, and the cecum (see Figure 1a). As expected, the abundant bacterial species differed significantly across the three organ sites with only a handful of species being common across the three communities (Figure 1b).

**Figure 1.**
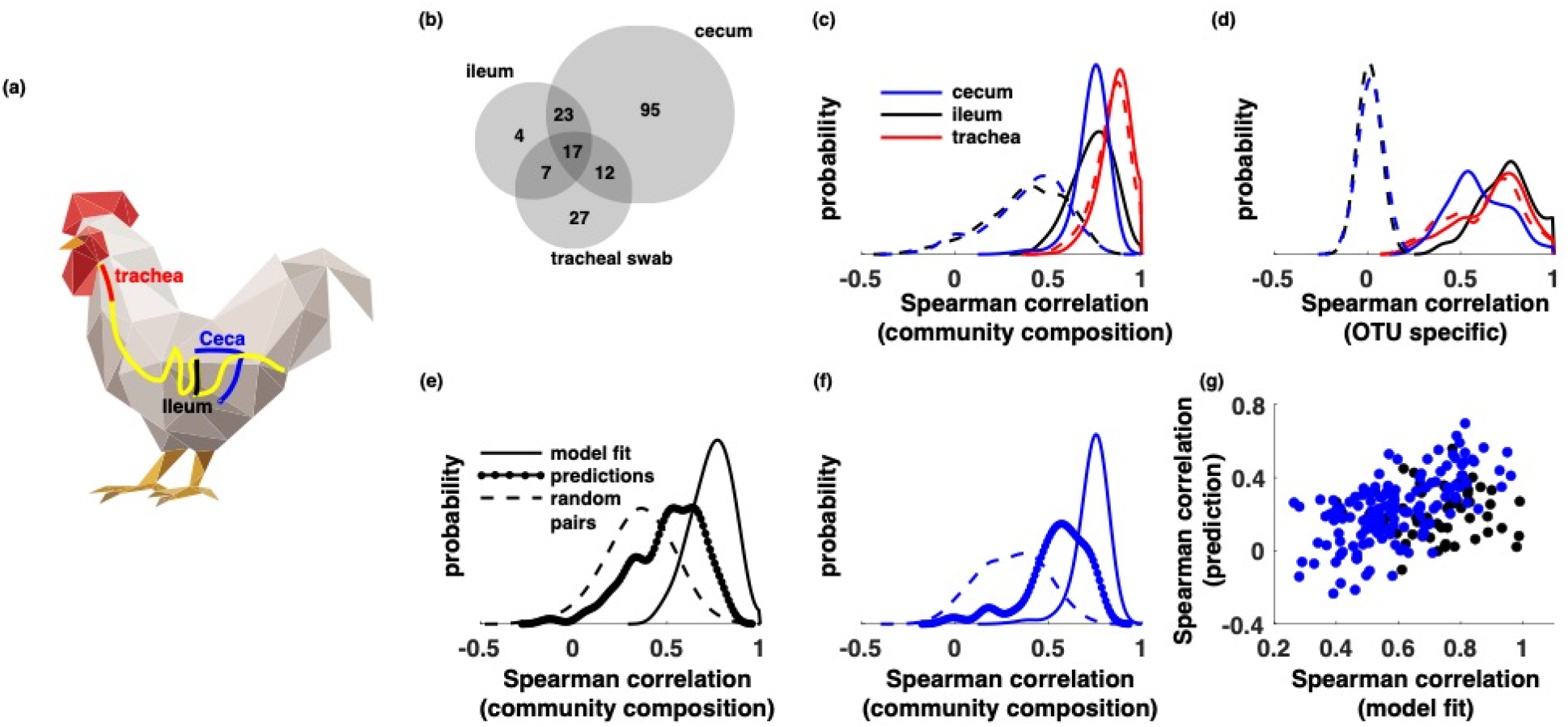
(a) Approximate location of the trachea, ileum, and ceca in a chicken’s digestive and respiratory tracts, **(b)** A venn diagram of the overlap between most abundant bacterial OTUs residing in the three organ sites. **(c)** The histogram of Spearman correlations between model fit community abundance profiles and measured community abundance profiles from samples collected from the cecum (blue), ileum (black), and the tracheal swabs (red). The same distribution when the communities were assigned to random hosts, thereby breaking their interdependence (dashed lines). (**d**) OTU-specific Spearman correlation coefficient (calculated across samples) between model fit and data for samples collected from the cecum (blue), ileum (black), and the tracheal swabs (red). The same distribution when the communities were assigned to random hosts, thereby breaking their interdependence (dashed lines). **(e)** The histogram of Spearman correlations between model fit community abundance profiles and measured community abundance profiles for the ileum (solid), the same distribution for randomly chosen pairs of communities (black), and for ileum community abundances predicted using communities in the trachea and the measured abundances in the ileum (dashed lines). (**f**) Same as (**e**) but for the cecum. (**g**) The OTU-specific Spearman correlation coefficient between model fit and data (x-axis) and the Spearman correlation coefficient between abundances predicted using the tracheal swab and data (y-axis) for cecum (blue) and the ileum (black). All histograms are smoothed using the MATLAB function *ksdensity* which uses a Gaussian kernel with a bandwidth of 0.05 and reflecting boundary conditions.

To identify the lower dimensional manifold that explains the covariation between these communities, we asked whether a small number of host-specific latent variables can explain the community composition at the three organ sites. Interestingly, even though the communities at the three different organ sites comprised different organisms, host-specific latent variables could explain community composition in the three organ sites in a host-specific manner (Figure 1c, d). The Spearman correlation coefficient between model fit community abundances and observed community abundances was *ρ* = 0.86 ± 0.07 for tracheal swabs, 0.74 ± 0.11 for the ileum, and 0.73 ± 0.08 for the cecum (Figure 1c). The model also captured OTU-specific abundances, the average correlations between model fit and measured OTU abundances were *ρ* = 0.67 ± 0.18 for tracheal swabs, 0.73 ± 0.14 for the ileum, and 0.59 ± 0.15 for the cecum respectively (Figure 2d). Notably, this agreement strongly relied on the communities being sampled from the same host. The model fit when hosts were assigned random communities performed significantly worse (dashed lines in Figure 1c, d). This suggests that host-specific variables were key determinants of compositions of host-associated communities.

**Figure 2.**
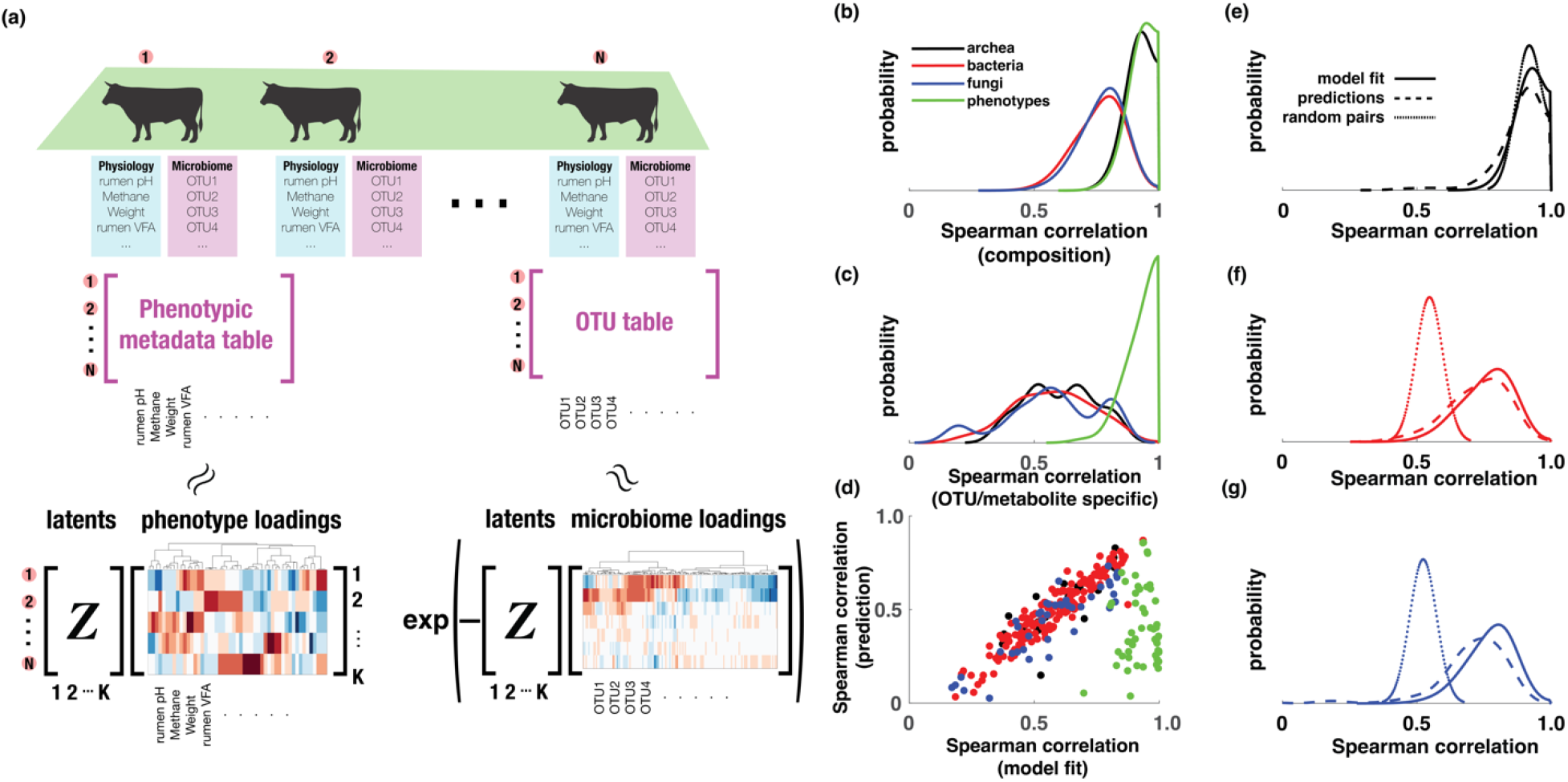
**(a) Schematic of SMbiot.** Individual hosts are characterized by multiple microbial ecosystems and host-specific phenotypic metadata. SMbiot models the covariation in the host-microbiome metacommunity using a common latent-space representation, **(b)** A histogram of Spearman correlation coefficients between measured community compositions and the corresponding model fits for archeal (black), bacterial (red), and fungal (blue) communities. Similar histogram is also shown for z-scores of host metadata (green) **(c)** A histogram of Spearman correlation coefficients between measured abundances of archeal (black), bacterial (red), and fungal OTUs (blue), as well as the correlation coefficient between measured host phenotypes and the corresponding SMbiot-based model fits (green), (d) A comparison of Spearman correlation coefficient between SMbiot-modeled OTU abundances and the measured OTU abundances (x-axis) and the Spearman correlation coefficient between SMbiot-predicted OTU abundances and the measured OTU abundances in a 80%/20% training testing split of the data. Data is shown for archeal (black), bacterial (red), and fungal OTUs (blue). Correlation comparison is also shown for individual host phenotypic data (green), (**e-g**) A histogram of Spearman correlations between model-fitted community compositions and measured community compositions (solid lines), Spearman correlations between predicted community compositions and measured community compositions (dashed lines) in a 80%/20% training testing split, and Spearman correlations between microbial communities from random pairs of hosts (dotted lines). Data is shown for archeal (black, e), bacterial (red, f), and fungal (blue, g) communities. All histograms are smoothed using the MATLAB function *ksdensity* which uses a Gaussian kernel with a bandwidth of 0.05 and reflecting boundary conditions.

While the host-specific latents were learned using all three communities, approximate information about latent space embedding can be obtained even with single communities. This has significant practical uses. Specifically, microbial communities from deep within the digestive tract, for example, those residing in the ileum and the cecum cannot be easily investigated without sacrificing the host organism. In contrast, microbial communities in the trachea can be investigated in live hosts. Therefore, it would be significantly useful if microbial community composition measured using tracheal swabs can predict communities in the ileum and the cecum. To test this, we randomly split the data into a 80%/20% training and testing split. First, we inferred the feature space representing the three communities using the training data. For the testing data, we inferred the host-specific latent space representation only using the community composition in the trachea and the feature space learned using the training data. Then, we predicted the community composition in the ileum and the cecum for the testing data. As shown in the dashed histograms in Figure 1e and 1f, predicted community composition agreed quite well with the measured composition both for ileal and cecal communities. The average community-wide Spearman correlation was *ρ* = 0.50 ± 0.13 for the ileum and 0.54 ± 0.17 for the cecum. These predictions were significantly stronger than the expected pairwise correlation between communities from the same organ site from random pairs of hosts in the testing data (black lines in Fig. 1e and 1f, Mann-Whitney U-test p values *p* = 7.4 × 10^−12^ and *p* = 4.2 × 10^−27^ for the ileal and the cecal communities). Notably, the accuracy of OTU-specific abundance prediction was largely governed by how well the model was able to capture the abundance of the OTU. As seen in Fig. 1g, cecal and ileal OTUs that were well fit using the latent space model were also well predicted using data collected on tracheal swabs.

The analysis performed here used a latent space with *K_C_* = 20 dimensions. However, the conclusions about the low-dimensional behavior were robust to changes in the latent-space dimension with higher dimensional descriptions leading to descriptions with somewhat higher accuracies (Supplementary Table 1).

### Host phenotypes explain host-specific latent variables

The analyses in Figure 1 conclusively show that host-associated microbial communities are controlled by a combination of host-specific variables. What dictates these variables? To investigate whether host phenotypes relate to the latent space description, we analyzed data collected by Wallace et al.^21^ which comprised separate measurements of compositions of three microbial kingdoms (bacteria, archaea, and fungi) in the bovine rumen as well as phenotypic characteristics of bovine hosts, including rumen metabolite levels, host physiology, and host blood protein levels (Figure 2a).

A typical host was characterized by ~ 275 attributes comprising relative abundances of ~ 225 highly abundant operational taxonomic units (OTUs) across the three kingdoms as well as ~ 50 phenotypic metadata. We trained a combined latent space model with *K_C_* = 5 – 20 latents that were shared across the descriptions of the three microbial kingdoms as well as the host metadata. The figures below are shown for *K_C_* = 20. Supplementary Tables 2–5 show that these conclusions do not depend strongly on the number of latent dimensions. We chose a shared latent approach to represent the bidirectionality between host-microbiome interactions.

First, similar to the chicken microbiomes, we examined whether SMbiot can accurately capture the observed variation in the microbiome(s) as well as the phenotypic data. In Figure 2b we show the histogram of the Spearman correlation coefficients between measured community compositions and community compositions reconstructed by the SMbiot-based latent space model. The histograms represent the three microbial kingdoms and the host metadata. SMbiot accurately captures the microbial community compositions with average Spearman correlations of *ρ* = 0.92 ± 0.04,0.76 ± 0.09, and 0.77 ± 0.08 for archaeal, bacterial, and fungal communities respectively. SMbiot also reproduced z-scores of individual metadata in hosts with an average Spearman correlation of *ρ* = 0.93 ± 0.04. In Figure 2c, we show that SMbiot accurately captures the variation in relative abundances of individual OTUs across hosts, as well as variation in individual host metadata across samples with average Spearman correlations of *ρ* = 0.60 ± 0.14,0.57 ± 0.16, and 0.57 ± 0.19 for archaeal, bacterial, and fungal OTUs respectively and average Spearman correlation of *ρ* = 0.93 ± 0.06 for host metadata features.

Similar to chicken microbiomes across multiple organ sites, the ability of SMbiot to accurately model the microbial communities and the host metadata depended on the covariation within metacommunity components. Indeed, when we trained an SMbiot-based model on metacommunities with randomly assigned microbiomes and host phenotypic information, SMbiot failed to model microbial compositions (Supplementary Figure 1).

In commercially raised cattle, an important feature that distinguishes host-microbial metacommunities is the farm identity. Cows in the same farm experience similar environments including food intake regimes and pathogen exposure. We wanted to test whether the inferred governing features (matrices corresponding to the metadata and the three microbial communities) were robust to inclusion of individual farms. To that end, for each farm ***F**,* we inferred a farm-specific SMbiot model (denoted by the superscript ***F***) and another model where we excluded the farm (denoted by the superscript *—**F***. Next, we inquired whether the corresponding governing features (denoted by and Θ^(***F***)^ and Θ^(**-*F***)^ could be projected onto each other using a matrix multiplication. If this is the case, then we can conclude that governing features corresponding to a specific modality (host metadata or one of the three microbial communities) are robust to inclusion/exclusion of specific farms. To do this, we inferred a rotation matrix ***B*** such that **Θ**^(***F***)^ ≈ BΘ^(***-F***)^ to assess similarity between individual feature matrices and evaluated the *L*_2_-norm *L*_2_ = |Θ^(***F***)^ – BΘ^(***-F***)^|. We compared the *L*_2_-norm above with the scenario where individual entries in the Θ^(***F***)^ matrices were randomized (denoted by 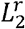). We found that in 14 out of the 20 projections considered (5 farms and 4 types of data), *L_2_* was statistically significantly lower than 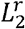 (Student’s t-test using 100 randomizations, Supplementary Table 6). This suggests that the learnt feature spaces represent true biological variability and are robust to inclusion/exclusion of farm-specific metacommunities. Interestingly, we find that the largest disagreement was between the fungal feature spaces. In fact, in 4 out of the 5 farms, the inferred fungal feature space with and without metacommunities from the specific farm could not be mapped onto each other.

Next, we inquired whether the feature spaces that embed individual samples allow us to predict microbial abundances and host metadata. To that end, we divided the samples using an 80% training/20% testing split. A SMbiot model was learned on the training data. For the test data, we identified the latent space description of individual samples using all but one piece of information about the hosts (either one of the three microbiomes or the host metadata). Then we evaluated whether we could predict the unseen data from the inferred latent space representation. As seen in Figure 2d, SMbiot can accurately predict abundances of OTUs for all three kingdoms of microbial life with average OTU-specific Spearman correlation coefficient *ρ* = 0.89 ± 0.08,0.73 ± 0.11, and 0.70 ± 0.16 for archaeal, bacterial, and fungal OTUs respectively. Similarly, SMbiot can also predict host metadata features just using the compositions of the host associated microbiomes. The average Spearman correlation coefficient of *ρ* = 0.41 across all metadata features with 50 out of the 53 features predicted with statistical significance with a 5% FDR correction (47 out or 53 features with a 1% FDR correction).

Interestingly, the microbial OTUs whose abundances were described well by SMbiot-based model were also the same OTUs whose abundances were predicted well. The Spearman correlation coefficient between measured OTU abundances and SMbiot-based fit was highly correlated with the Spearman correlation coefficient between measured OTU abundances and SMbiot-based predictions. However, this trend was not observed in the metadata. Virtually all metadata components were well described by the SMbiot-based model. In contrast, only 16 out of the 53 of the total metadata components could be predicted with a Spearman correlation coefficient greater than *ρ* = 0.5 using the compositions of the three microbiomes. The host metadata that were well-predicted were associated directly with the rumen or with hosts’ diet. In contrast, the metadata that were less accurately predicted were associated with phenotypes unrelated to the microbiome(s), for example, blood haptoglobulin levels and cholesterol levels (Supplementary Table 7). This shows that SMbiot-based predictions can potentially be used to identify direct or causal variations in the metacommunity.

Similar to predicting abundances of individual OTUs across different samples, SMbiot can also predict compositions of entire microbial communities (Figure 2e-g) with an accuracy that is comparable to SMbiot-based model fits. Notably, this prediction accuracy was not due to similarity between community compositions across hosts. Indeed, the Spearman correlation between microbial communities from random pairs of hosts was significantly smaller than SMbiot-based predictions.

These two analyses convincingly show that variation in host associated microbial communities can be explained using a small number of governing variables and that these variables are related to the host’s phenotypic states.

### SMbiot highlights the context-specificity of microbiome-phenotype associations

Covariations within the host-microbiome metacommunity are context-dependent, that is, the correlation between a bacterial species and a host phenotype can be altered by the presence/absence of other bacteria in the ecosystem as well as environmental factors. SMbiot allows us to systematically evaluate this context dependence of the covariation within metacommunities. For concreteness, let us consider a SMbiot model to describe abundances of host associated microbial species and host phenotypes. For any point ***z*** in the latent space (for example, a point representing a host *S* used for training the model), we can define the matrix of interactions **Δ**(***z***) that relates a small change in microbial abundances *δ_q_o__* to small changes in host phenotypes *δ_μ_p__*,

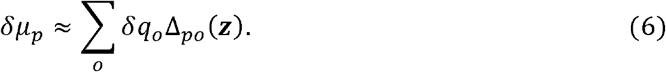

SMbiot allows us to analytically obtain the functional dependence of the interaction matrix on the latent space location (**Supplementary Material**). In other words, SMbiot predicts that the effect of the same change in the microbiome can lead to a different change in host phenotypes for different hosts.

To demonstrate the host-specific nature of the sensitivity matrix, we used data from the Inflammatory Bowel Disease Multiomics Database^22^. Here, several paired human fecal microbiome and gut metabolome samples were collected from the same individual for multiple individuals. We first trained a SMbiot model using paired microbial composition and fecal metabolomics data from a single randomly chosen sample per individual. As with the bovine and the chicken metacommunities, a SMbiot model with a very small number of latents was able to fit the host-specific covariation between microbial species abundances and host gut metabolomics (Supplementary Figure 2). Next, we used the additional microbiome/metabolomics samples from the same individuals treated as small perturbations on the training samples. We then estimated the predicted changes in the metabolome given a change in the microbiome using the subject-specific interaction matrix. The host specific interaction matrix could accurately predict the direction of large (>2 standard deviations) abundance changes of individual metabolites (Fig. 3a) and the magnitude of the change of individual metabolites (e.g. Butyrate, Fig. 3b). Importantly, these predictions were context dependent. Using an interaction matrix from a randomly chosen host led to significantly lower prediction accuracy (Fig. 3a,b, dashed lines). Importantly, the extent of the microbiome differences between reference and perturbation samples used for making predictions are identical, with only the interaction matrix being changed.

**Figure 3.**
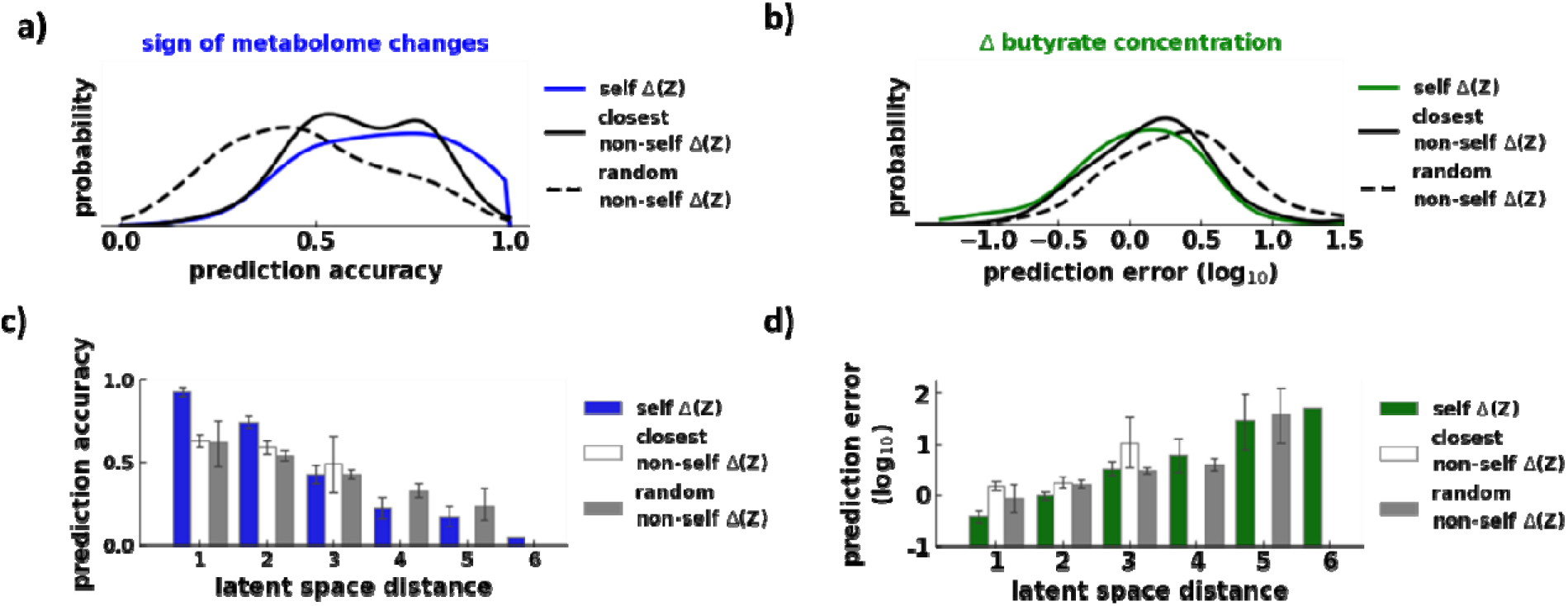
(a) Histograms for accuracy values for predictions of the direction of metabolite changes based on the interaction matrix associated with a sample from the same individual on which the changes were measured (solid blue), an interaction matrix from a randomly selected individual (dashed blac), or the interaction matrix from a different individual closest in the latent space to the reference sample (solid black). Accuracy was calculated for changes > 2 s.d. (b) Histograms for the prediction error in the magnitude butyrate abundance changes using an interaction matrix from the same individual on which butyrate abundances were measured (solid green), a randomly chosen different individual (dashed black), or a different individual closest in the latent space (solid black). (c) The relationship between accuracy of predictions of the direction of metabolite changes and the distance in the latent space between the sensitivity matrix used and the reference sample on which changes were measured. (d) Like c but for the error in the prediction of butyrate abundance changes.

This context-specificity is largely driven by the fact that samples from the same individual tend to be closer in the latent space than samples from different individuals. For example, using an interaction matrix from a different host individual that is closest in the latent space to the reference host (Figure 3a,b) led to slightly lower but similar prediction accuracies as using the sensitivity matrix from the reference individual. When we binned predictions by the distance in the latent space between reference samples and the sensitivity matrix used to predict metabolome changes, the accuracy of phenotype response predictions decreased for larger distances in the latent space (Figure 3 c,d). The dependency of prediction accuracy on the sensitivity matrix used underscores the context-specificity of host-microbiome interactions and the ability of SMbiot to capture this context-specificity.

### SMbiot predicts the effect of antibiotic treatment on the chicken ceca metabolome

To validate the predictive ability and to illustrate a typical practical application of SMbiot, we used recently published metabolomics and bacterial abundance data from chickens^23^. The data was originally used to assess the impact of different antibiotics on the composition of the cecal microbiome and metabolome. Although microbial profiles were determined for birds receiving no antibiotics, avilamycin, or narasin in the diet, metabolomics data were only published for control birds and birds fed with narasin. We trained a SMbiot model on the data from control and narasin birds and used it to predict the abundances of ~450 metabolites in birds fed with avilamycin from the reported microbial abundances. For validation, we experimentally obtained metabolite abundances for samples from the same individuals.

Similar to the above 3 metacommunities, the host specific covariation between cecal microbiomes and metabolomes of chickens in the training data was accurately captured by a shared latent model (Supplementary Figure 4). Importantly, the correlation between normalized metabolite abundances and corresponding SMbiot predictions was significantly higher than the similarity between random pairs of birds in the training data (**Fig. 4a**, median p-value=0.03). At the level of individual metabolites, ~ 75% of compounds showed a positive correlation between predictions and experimental measurements, and this correlation was significant for about 10% of compounds (*r* > 0.51) across the n=15 birds in the testing set (**Fig. 4b**). In addition, the average change in metabolite abundances between control birds and avilamycin treated birds correlated significantly between SMbiot predictions and measured data (**Fig. 4c**, Spearman *r* = 0.49, *p* < 10^−30^); with the predicted direction of the change in metabolite abundances between the two treatments agreeing in ~ 70% of cases (binomial *p* < 10^−6^). The results show that SMbiot can be used to extend previously available multi-omics datasets, to impute missing experimental data, and to generate and support hypotheses about the association of microbes with host phenotypes. For instance, Plata et al. reported lower abundances of amino acids and long-chain fatty acids in the cecum of birds treated with narasin^23^, SMbiot predictions accurately showed similar patterns for 17/20 amino acids and 14/14 long-chain fatty acids with avilamycin (**Fig. 4d**). Suggesting that avilamycin has similar effects on the cecal metabolome as narasin.

**Figure 4.**
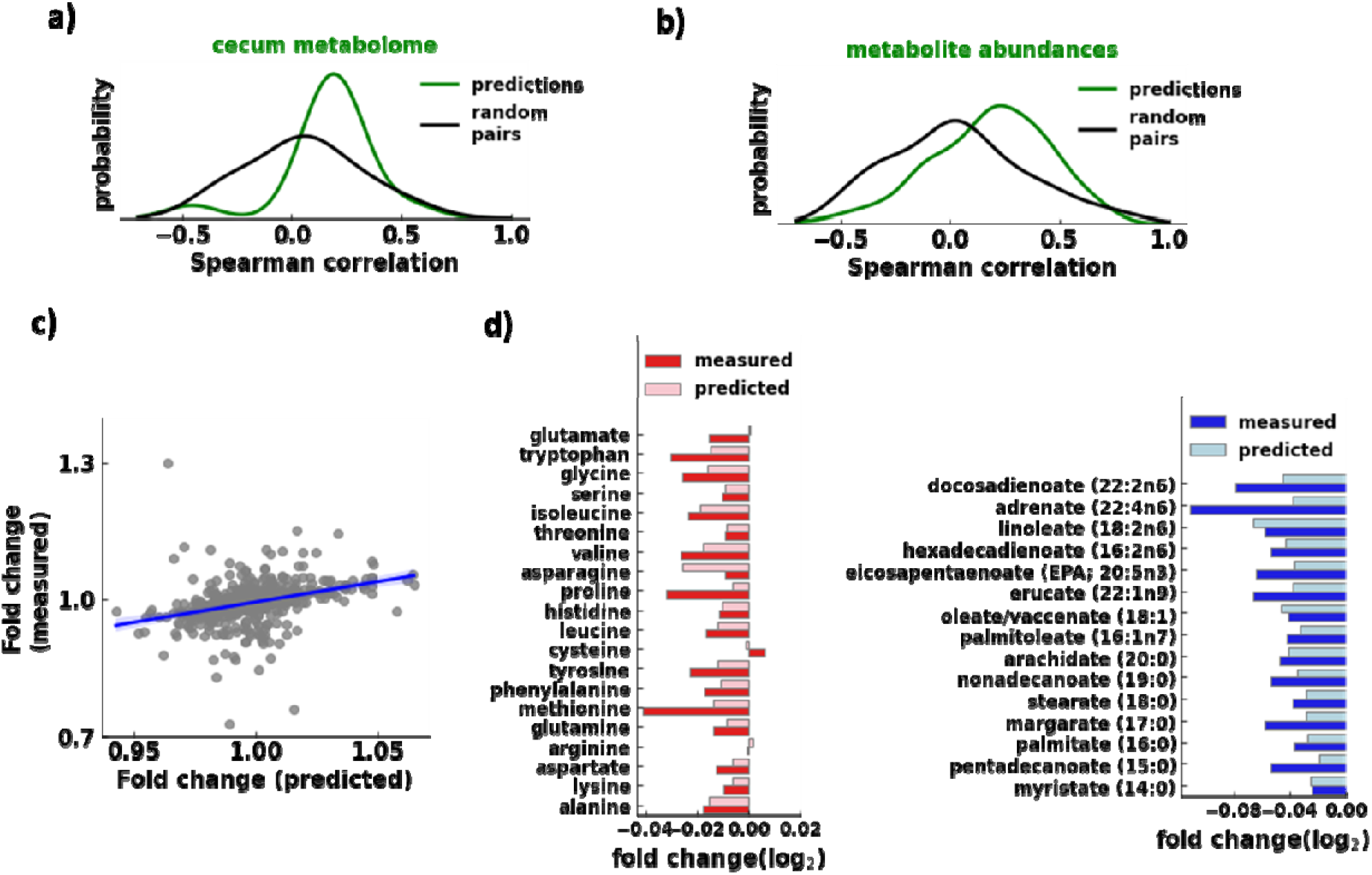
**(a)** The histogram of Spearman correlations between SMbiot predicted metabolite profiles in avilamycin treated birds, and experimentally measured metabolite profiles (green). The black line shows the distribution of metabolite profile correlations between random pairs of samples, (b) The histogram of Spearman correlations between SMbiot abundance predictions for individual metabolites and experimental measurements (green) across samples (n=l5). The black line shows the distribution of correlations between random pairs of metabolites, (c) SMbiot predicts the direction and relative magnitude of metabolite abundance changes in avilamycin treated birds compared to control birds (Spearman r= 0.52, p<1e-30). The Blue line shows a linear fit and 95% confidence interval, (d) SMbiot accurately predicts a decrease in the abundance of amino acids and long chain fatty acids in the cecum of birds treated with avilamycin compared to controls.

## Conclusions

The fate of microbial species colonizing animal host organs are determined by the physiological state of the host, organ-specific effects including nutrient availability and host immune system presence, as well as competitive and cooperative interactions with each other within and across kingdoms of life.

Due to these many interactions, it is reasonable to expect that compositions of individual microbial communities do not vary independently of each other but rather reside on a manifold of a much smaller dimension. We presented SMbiot, a shared-latent based approach to identify smaller dimensional description of covariation within microbial communities associated with the same host using host-specific latent variables. We showed that these latent variables tightly associate with hosts’ phenotypic states. We also showed that SMbiot allows us to predict unmeasured data based on partial information. Importantly, SMbiot could also quantify the context-specificity of host microbiome interactions. These analyses conclusively show that covariation in host physiological states and the associated microbiome(s) can be understood using a small number of controlling variables. Going forward, we believe that SMbiot will be an important computational tool to study host microbiome interactions.

## Acknowledgement

PD acknowledges the support of NIH grant R35GM142547

Code: https://github.com/dixitpd/SMbiot/

## Methods

### SMbiot:A Shared Latent Model for Microbiomes and their Hosts

An animal host and its associated microbial ecosystem(s) can be described by several quantifiers such as the blood (or fecal) metabolome of the animal, animal physiology (weight, diet intake, etc.), and microbial composition (bacterial, archaeal, fungal, etc.) at various host sites (skin, intestine, etc.). **SMbiot** (pronounced SIM BY OT), A **S**hared Latent **M**odel for Micro**bio**mes and their Hos**t**s is an integrated approach that simultaneously models these quantitative features of the host and the associated microbial ecosystem(s). For concreteness, we present our development using the case when data on animal physiology and bacterial microbiome at one site are available for several hosts. Specific generalizations to other types of data will be discussed as well.

Let us consider that we have measured *P* phyiological metadata features and abundances of *0* operational taxonomic units (OTUs) in S hosts (samples). We denote the individual metadata features as *m_sp_* (*s* ∈ [1, *S*], *p* ∈ [1, *P*]) and individual OTU read counts as *n_so_* (*s* ∈ [1, *S*],*o* ∈ [1, *O*]. We seek a reduced dimensional representation of the metadata and the microbiome wherein some of the latent variables (low dimensional features) are shared between the metadata and the microbiome. To that end, we model the metadata (assumed to be Z-transformed) as a multivariate Gaussian distribution:

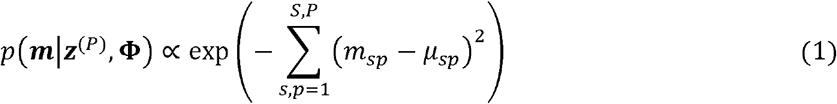

where

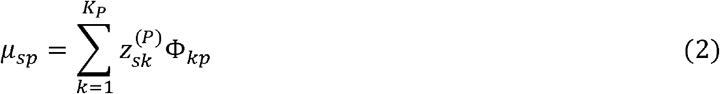

is the low-rank approximation. We choose *K_P_* ≪ *S, P* to impose the low rank. In Eq. (2), ***z***^(*P*)^ are collectively the latents that describe the metadata and **Φ** is the matrix of physiology-related features. We note that other model distribution choices (for example, exponential or log-normal distribution) are possible as well.

We model the microbiome abundances (for example, OTU read counts) using a multinomial distribution,

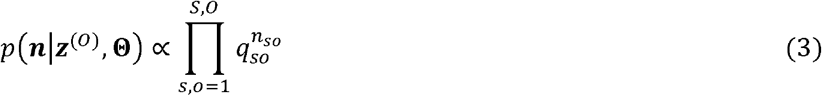

where

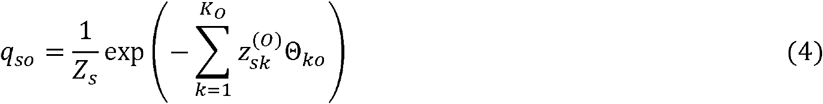

is the reduced-dimensional description of the microbiome (cite). In Eq (4) ***Z***^(O)^ are collectively the latents that describe the microbiome and Θ is the matrix of microbiome features. We have previously shown that this Gibbs-Boltzmann form can approximate microbial abundances to a high degree of accuracy using a very small number of latents^1^.

A salient feature of SMbiot is that we obtain an integrated description of the host and the microbiome by imposing that *K_C_ ≤ K_O_, K_P_* latents are shared between the latent space approximation of the microbiome and the metadata. Concretely, we enforce

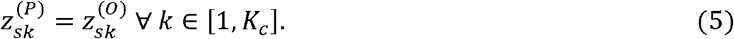

We infer the model parameters {***z***^(*P*)^, ***z***^(*O*)^, **Φ, Θ**} by maximizing the combined log-likelhood of the metadata and the microbiome. We write the weighted log-likelihood

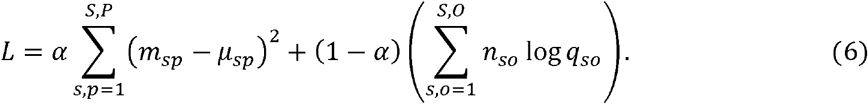

In Eq. (6), the constant 0 < *α* < 1 dictates the relative importance of the metadata or the microbiome in the shared latent dimensionality reduction approach; *α* ~ 1 prioratizes the metadata over the microbiome and vice versa for *α* ~ 0. The normalization constants in the log likelihood that do not depend on model parameters are omitted.

The gradients of the likelihood with respect to the model parameters; the latents ***z***^(*P*)^,***z***^(*O*)^, and the features Φ and Θ can be derived analytically. We use these gradients to infer the model parameters in a maximum likelihood approach. We have

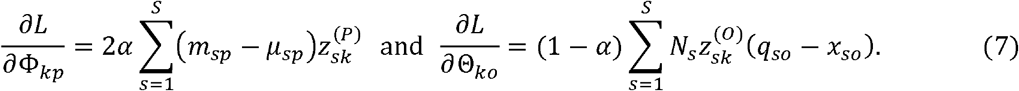

where *N_s_* = ∑*_o_ n_so_* is the total read count of the sample and *x_so_= n_so_/N_s_* are relative OTU abundances. Similarly, we have

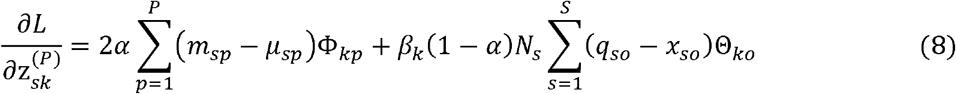

and

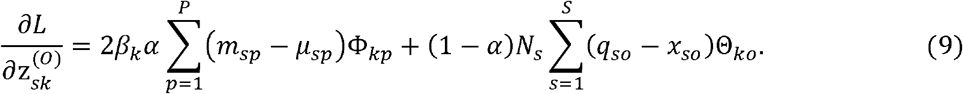

In Eq. (8) and (9), *β_k_=* 1 if *k < K_c_* and zero otherwise. We identify the model parameters using simple gradient ascent with a constant learning rate, ranging between *η =* 10^−3^ to *η =* 5 × 10^−3^.

In this general shared latents formalism, an animal host and the associated ecosystem can be represented using other types of data as well. For example, we may have data available for simultaneous microbial composition at several host sites such as oral swabs, intestine, and feces, in addition to information about the hosts’ physiology. In this case, we will amend the log-likelihood function (Eq. 6) with several multinomial distributions. If several types of orthogonal metadata are available, for example, fecal and serum metabolomics, we will amend the log-likelihood function with an additional Gaussian distribution. Below, we show how to extend this formalism for specific datasets.

Finally, we note that the latent space and the governing axes (matrices **Θ** and **Φ**) inferred using maximization of Eq. 5 are not uniquely determined but belong to a linear space whose dimension equal to the dimension of the latent space. Specifically, if we denote by **Θ**^(**1**)^ and **Θ**^(**2**)^ (and **Φ**^(**1**)^ and **Φ**^(**2**)^ the SMbiot-learnt matrices from two different initializations, there exists a *K × K* invertible matrix *B* such that Θ^(**1**)^ =*BΘ*^(**2**)^ and **Φ**^(**1**)^ =*B**Φ**^(**2**)^*. Notably, the inverse of this matrix will also transform the latent space: ***Z***^(**1**)^= B^−1^***Z***^(**2**)^.

### Using SMbiot to make predictions from partial data

#### SMbiot predicts the microbiome composition given physiological metadata and vice versa

The shared latent framework of SMbiot allows us to predict the microbiome of a host given the metadata m^host^λ Using the metadata vector, we can obtain the 1 ×*K_P_* vector of metadata latents z^(P,host)^ by maximizing the likelihood in Eq. (1). For Gaussian likelihood, this can be done analytically.

We have

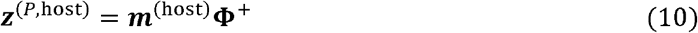

where Φ^+^ is the Penrose-Moore pseudoinverse of the feature matrix Φ. When analytical estimates are not possible, we can infer z(^*p*,host^) using maximum likelihood.

Only *some* of these inferred latents ***z***(^p,host^) are shared with the microbiome which we denote by ***z***^(p,host)^. We model the remaining microbiome latents using a multivariate Gaussian distribution. Here too, other approaches to guess the unconstrained microbiome latents are possible as well. We assume that the microbiome latents are distributed according to a *K_o_* dimensional multivariate Gaussian distribution. The means 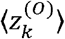 and the covariance matrix elements ∑_*kl*_ of this distribution can be estimated from the training samples:

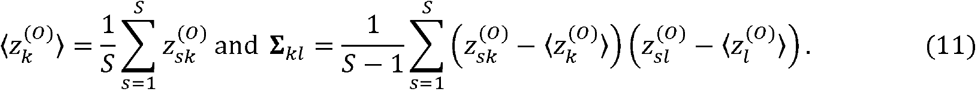

The conditional distribution of the unconstrained latents given the shared latents is also a Gaussian distribution albeit with modified mean values and covariances (cite). The conditional means of the unconstrained latents are given by

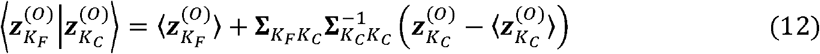

and covariances are given by

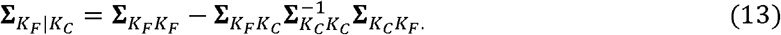

In Eq. (12) and Eq. (13), a subscript *K_F_* denotes the collective indices of the free microbiome latents and *K_C_* denotes the collective indices of the microbiome latents shared with the metadata. The matrices (vectors) with the corresponding subscripts denote block matrices (vectors) with the corresponding indices. This way, the metadata and the multivariate Gaussian model gives us all microbiome latents. Once the latents are known, the microbiome composition can be modeled using the Gibbs-Boltzmann distribution Eq. (4). The same approach can be taken when predicting the metadata from the microbiome. Specifically, microbiome-specific latents are first inferred using only the microbiome composition and the remaining metadata components can be filled in using a multivariate Gaussian model. The Gaussian approximation is not needed if no “free” latents are employed.

#### Elucidating context-specificity of perturbations using SMbiot

Let us consider that we have collected data on the microbiome composition and the physiological metadata of some hosts and that we have modeled these data using the SMbiot approach. We assume that there are a total of *0* microbial species and *M* metadata. We assume that there are *K_tot_* latents. We assume that the latents are organized as follows: latents *K* = 1 to *K = K_q_* are solely coupled to the microbiome. Latents *K = K_q_* + 1 to *K = K_q_ + K_c_* are coupled to both the microbiome and the metadata, finally, latents *K = K_q_ + K_c_* + 1 to *K = K_q_ + K_c_ + K_m_= K_tot_* solely couple to the metadata. For any point *z* in the latent space (represented by a 1 ×*K_tot_* vector), we write the model approximations:

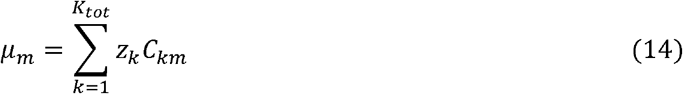

and

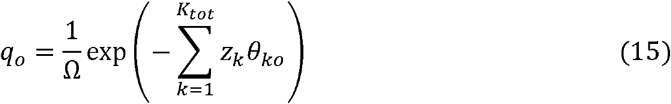

where *μ_m_* is the *m*^th^ metadata and *q*_0_ is the relative abundance of the *o*^th^ OTU. We note that given how the latents are organized, some rows in the *C* and the *θ* matrices will be zero. Specifically, *C_km_* = 0 ∀ *k* ∈ [1, *K_q_*] and *θ_ko_*= 0 ∀ *k* ∈ [*K_q_* + *K_c_* + 1, *K_tot_*], For any point *z* in the latent space, we can consider a small perturbation in the latents *dz* around the original vector of latents *z* and inquire how does this change affect the microbiome and the metadata. We have

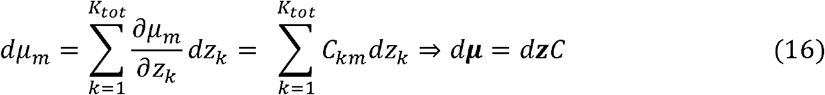

where *dμ* is a 1 ×*M* vector of changes in metadata and *dz* is a 1 × *K_tot_* vector of changes in the latents. Similarly, for the change in the microbiome, we can write

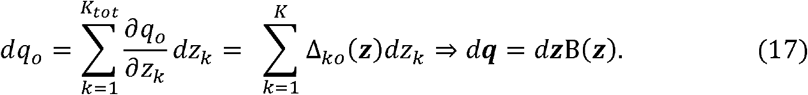

We will derive the functional form of the sensitivity matrix Δ(**z**) below. We note that the sensitivity matrix B(**z**) is a function of **z**, the latent space location. From Eq. 3 and 4, we can write:

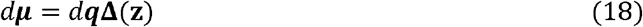

where **Δ**(***z***) = **Γ**(**z**)**C** and **Γ**(**z**) = (B(***z***))^+^ is the pseudoinverse of the matrix B(***z***). Next we derive the functional form of B(***z***). We first note that

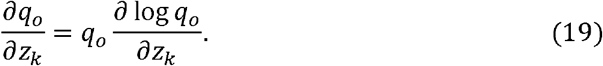

We have,

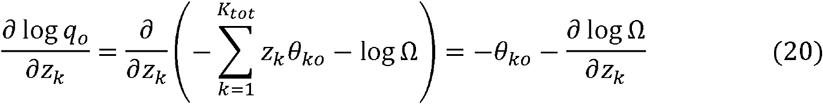

where

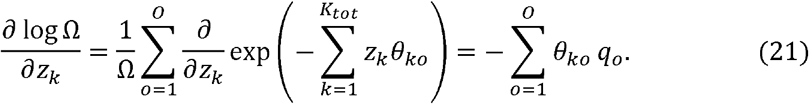

Therefore

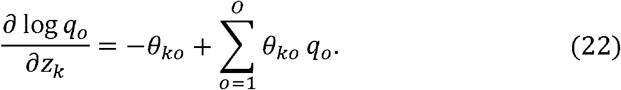

The sensitivity with respect to absolute changes is given by

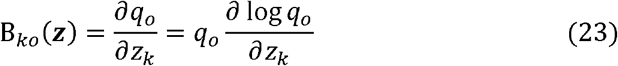

We obtain the **z**-dependent interaction matrix by substituting Eq. 23 in Eq. 18.

#### Descriptions of the datasets

Four datasets were used in our study. These datasets spanned multiple animal hosts, kingdoms of microbial life, host organ sites, and physiological data types. We used data collected on (1) microbial compositions across three kingdoms of microbial life (archaea, bacteria, and fungi) in the bovine rumen as well as physiological information about the bovine host^2^, (2) bacterial ecosystems in three different organ sites in a chicken^3^ (tracheal swabs, ileum, and cecum), (3) fecal bacterial ecosystem composition and fecal metabolomics in humans^4^, and (4) chicken cecal bacterial ecosystem composition and cecal metabolomics^5^. Below, we provide details of individual data sets

#### Three kingdoms of life in the bovine rumen and host physiology

The data on rumen microbiomes and host physiology of Holstein cows was downloaded from the paper by Wallace et al. (cite). We only included the samples that had a total read count of at least 2000 reads for each of the three microbial kingdoms. We only included the physiological metadata features that were measured in more than 50 hosts. For each kingdom, we included OTUs whose average relative abundance was larger than 0.1%. This cutoff was based on our previous work where we had shown that OTU abundance variation below 0.1% is likely to be dominated by technical noise^6^. In the end, we had data on 787 cows, described by relative abundances of 26 archaeal OTUs, 157 bacterial OTUs, 41 fungal OTUs, and 53 physiological metadata features. For each kingdom of microbial life, one of the OTUs represented the combined abundance of all OTUs whose average abundance was below 0.1%. We trained an SMbiot-based latent space model for the hosts using *K_C_* = 5 – 20 latents. All latents were shared across all three kingdoms of microbial life and the physiology. Given that we had three ecosystems, we modified the likelihood function presented above. The likelihood function used for model fitting was given by:

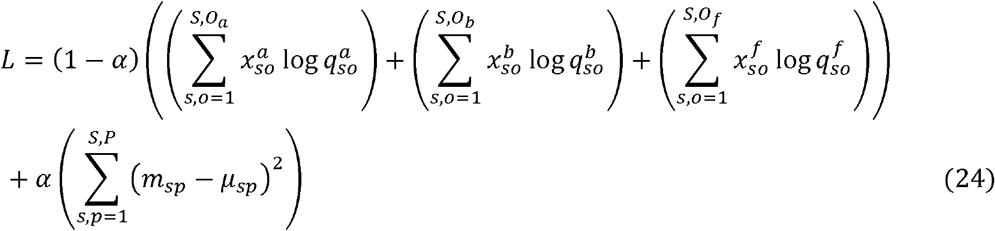

In Eq. X, we chose *α* = 0.1. The subscripts and the superscripts a, b, and f denote the archaeal, bacterial, and fungal ecosystems. The relative abundances are denoted by xs and the *ąs* are the corresponding model fits. The *ąs* were modeled using the model in Eq. 4. The ms denote z-scored physiological metadata features and the corresponding *μs* are model fits.

#### Chicken microbiomes

The data on chicken microbiomes from tracheal swabs, ileum, and cecum was downloaded from the paper by Johnson et al.^3^. We only included the hosts with samples that had a total read count of at least 1000 reads for each of the three organ sites. For each organ site, we only included OTUs whose average relative abundance in that organ site across all chickens was larger than 0.1%. In the end, we had a total of 425 host samples. Each host sample was described by three microbial ecologies comprising 63, 51, and 147 OTUs in the tracheal swabs, ileum, and cecum respectively. In each organ site, one of the OTUs represented the combined abundance of all OTUs whose average abundance was below 0.1%. We trained an SMbiot-based latent space model for the hosts using *K_c_ =* 5 – 20 latents. All latents were shared across all three organ sites. Given that we had three bacterial ecosystems and no physiological metadata, we modified the likelihood function presented above. The likelihood function used for model fitting was given by:

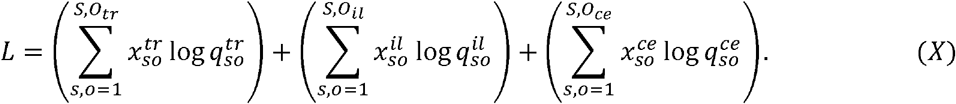

In Eq. X, the superscripts (and subscripts) tr, ol, and ce represent trachea, ileum, and cecum respectively. The relative abundances are denoted by xs and the *qs* are the corresponding model fits. The *qs* were modeled using the model in Eq. 4.

#### Human fecal metabolome and microbiome

Metagenomics-derived genus-level microbial abundances and untargeted metabolomics data were obtained from the Inflammatory Bowel Disease Multiomics database^4^. We considered 380 samples that had both metagenomics and metabolomics data available and an associated subject ID (obtained from the Sequence Read Archive). Only samples from subjects with 2 or more samples were considered for analysis. As above, genera with average relative abundances across samples lower than 0.1% were aggregated into a single OTU, yielding a total of 43(+1) bacterial genera analyzed. We considered for analysis 239 named metabolites with measured intensity greater than zero across all 239 samples. The intensities of mass-spectrometry peaks identified as the same metabolite were summed before analysis. Metabolite data were log10 transformed *m*→ log_10_(*m* + 1) before fitting the SMbiot model.

#### Chicken cecal microbiome and metabolomics

16S rDNA Amplicon Sequence Variant (ASV) abundances for control, narasin-treated and avilamycin-treated birds, as well as metabolite abundances for control and narasin-treated birds were obtained from Plata et al.^5^. N=15 samples per treatment. For training the model, ASVs with average relative abundances across samples lower than 0.1% were aggregated into a single OTU, resulting in 129(+1) ASVs analyzed. The values for metabolites with measured intensity greater than zero across all 30 training samples were loglO transformed before fitting the model. For validation of SMbiot predictions, untargeted metabolomics of the 15 samples from avilamycin-treated birds was carried via Ultrahigh Performance Liquid Chromatography-Tandem Mass Spectroscopy (UPLC-MS/MS) as described in Plata et al. The animal experiments and procedures were approved as described in Plata et al.

## Supplementary Figures

**Supplementary Figure 1.**
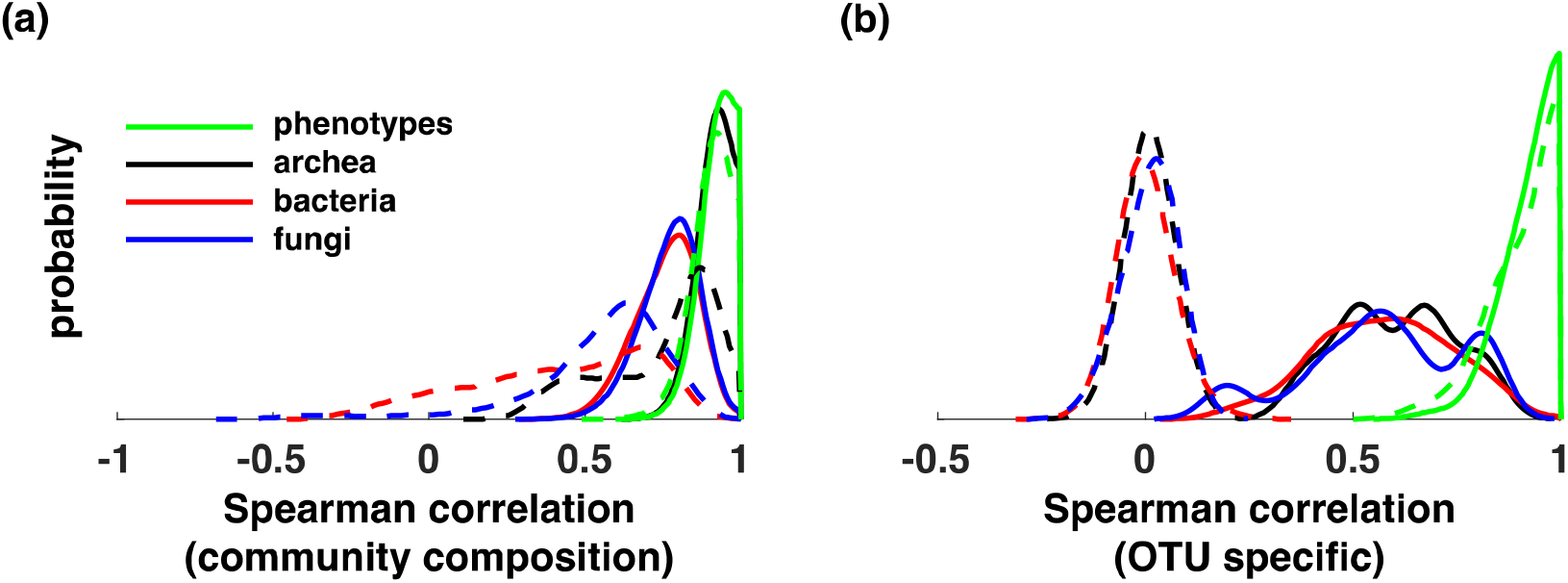
**(a)** Distribution of Spearman correlation coefficient calculated between model fits of community compositions (and z-scores of host phenotypic metadata) and the corresponding data. Solid lines represent an SMbiot model fit to collected data and dashed lines represent SMBiot model fit to randomized data wherein microbial communities were randomly assigned to hosts, (b) Same as in (a) but for OTU-specific correlations. components were used to model the data.

**Supplementary figure 2.**
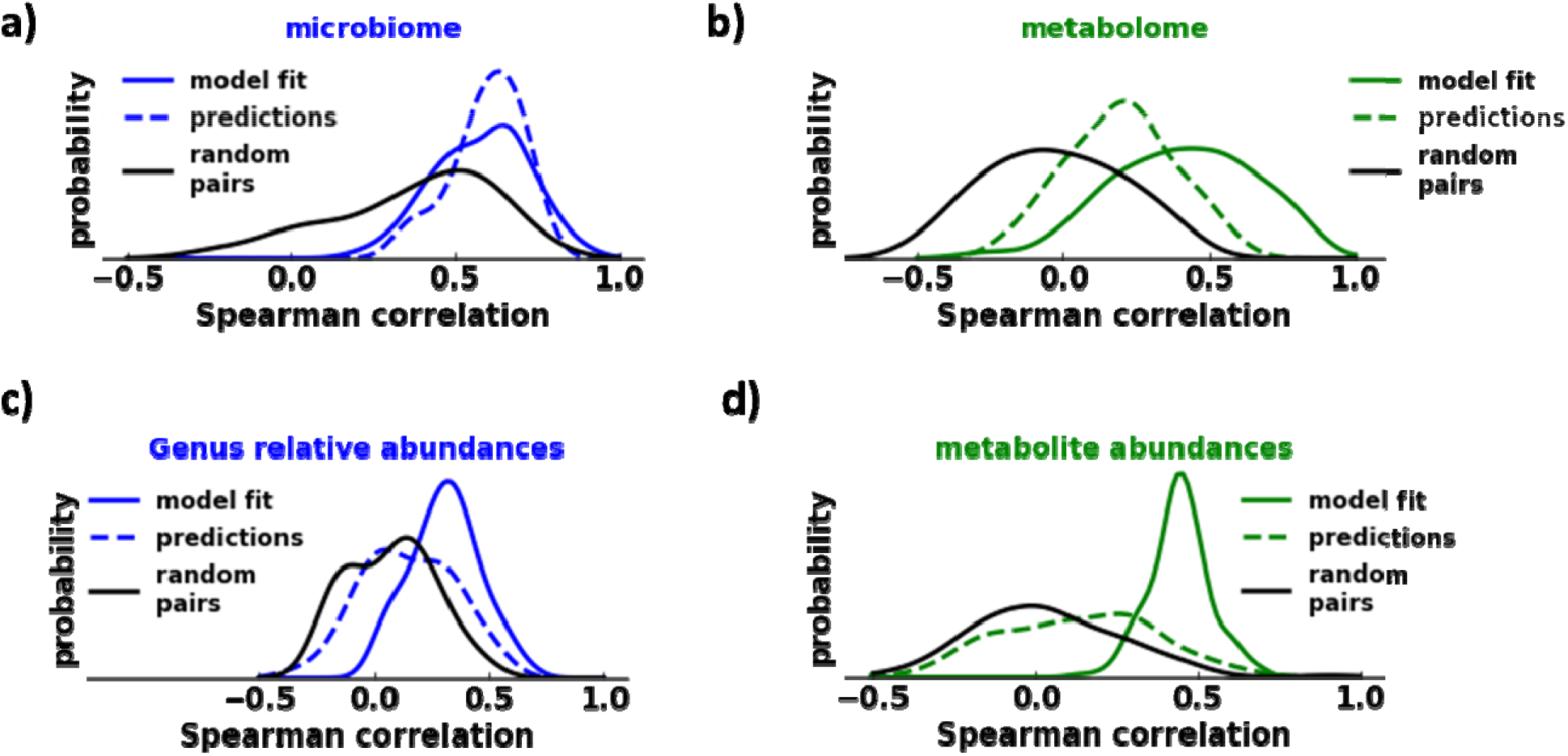
Model fit to fecal microbial and metabolite abundances in human samples, (a) Histograms of Spearman correlations between measured genus level microbiome profiles within a sample and SMbiot model fits (solid blue) or microbial profile predictions (dashed blue). The black histogram shows the distribution of correlations for random pairs of samples, (b) Like a, but for prediction of metabolite profiles within a sample, (c) Histograms of Spearman correlations between the relative abundance of individual genera across samples and SMbiot model fits (solid blue) or SMbiot predictions (dashed blue). The distribution of correlations between random pairs of OTUs across samples in shown in black, (d) Like c, but for the abundance of individual metabolites across samples. We have used latents shared between the medata and the microbiome and two extra latents to fit the microbiome alone.

**Supplementary Figure 3.**
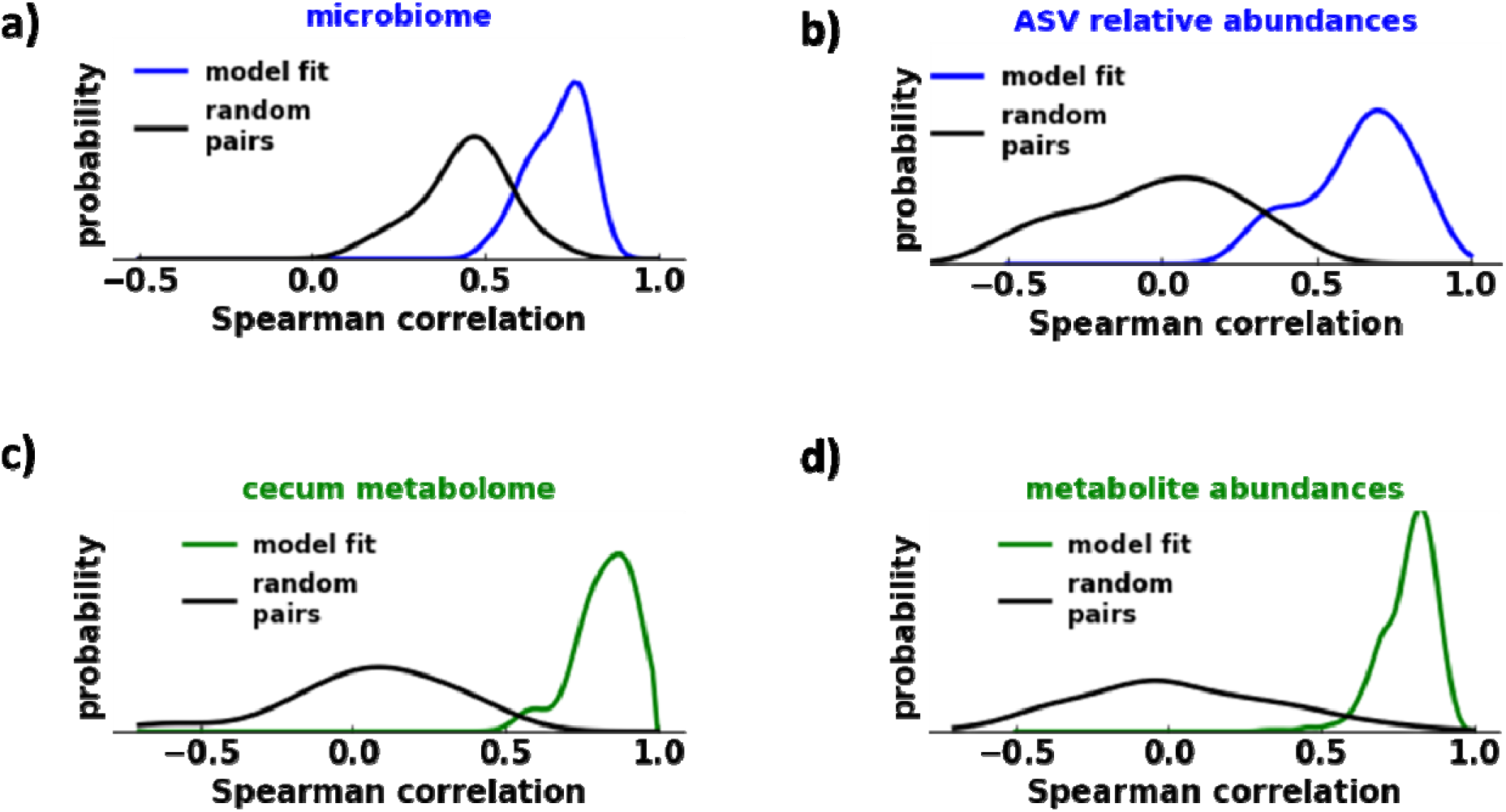
Model fit to cecal microbial and metabolite abundances in poultry, **(a)**The histogram of Spearman correlations between measured ASV abundance profiles and SMbiot model fits (blue). The black line shows the distribution for random pairs of samples, (b) The histogram of Spearman correlations between the relative abundances of individual ASVs across samples (blue). The black line shows the distribution for random pairs of ASVs. (c) Like a, but for metabolite abundance profiles, (d) Like b, but for individual metabolites across samples. We have used latents shared between the metadata and the microbiome and two extra latents to fit the microbiome alone.

## Supplementary Tables

**Supplementary Table 1.** Average Spearman correlation (and standard deviation) as a function of number of latents for microbial communities across the three organs.

| Tracheal Swabs |  | # of latents | Community | OTUs |
| --- | --- | --- | --- | --- |
| | | 5 | $0.79 \pm 0.08$ | $0.45 \pm 0.15$ |
| | | 10 | $0.82 \pm 0.08$ | $0.58 \pm 0.16$ |
| | | 15 | $0.84 \pm 0.08$ | $0.62 \pm 0.17$ |

| Ileum | # of latents | Community | OTUs |
| --- | --- | --- | --- |
| | 5 | $0.65 \pm 0.11$ | $0.59 \pm 0.15$ |
| | 10 | $0.69 \pm 0.10$ | $0.67 \pm 0.15$ |
| | 15 | $0.71 \pm 0.11$ | $0.70 \pm 0.15$ |
| Cecum | # of latents | Community | OTUs |
| | 5 | $0.61 \pm 0.10$ | $0.36 \pm 0.20$ |
| | 10 | $0.65 \pm 0.09$ | $0.47 \pm 0.17$ |
| | 15 | $0.70 \pm 0.09$ | $0.54 \pm 0.16$ |

**Supplementary Table 2.** Average Spearman correlation (and standard deviation) as a function of number of latents. The parantheses “model” indicates the correlation coefficient was calculated between data and corresponding model fit. The parantheses “predictions” indicates that the correlation coefficient was calculated between testing data and model predictions (using a 80%/20% training/testing split). “Community” correlation coefficients measure the accuracy of reconstruction of community composition. “OTUs” correlation coefficients measure the accuracy of OTU abundance reconstruction across host samples. The numbers are shown for the archeal community.

| Archea | # of latents | Community (model) | OTUs (model) | Community (predictions) | OTUs (predictions) |
| --- | --- | --- | --- | --- | --- |
| | 5 | $0.82 \pm 0.07$ | $0.40 \pm 0.21$ | $0.82 \pm 0.07$ | $0.37 \pm 0.21$ |
| | 10 | $0.88 \pm 0.07$ | $0.54 \pm 0.15$ | $0.87 \pm 0.07$ | $0.51 \pm 0.15$ |
| | 15 | $0.91 \pm 0.04$ | $0.58 \pm 0.14$ | $0.87 \pm 0.11$ | $0.51 \pm 0.15$ |
| | 20 | $0.92 \pm 0.04$ | $0.60 \pm 0.14$ | $0.89 \pm 0.08$ | $0.52 \pm 0.16$ |

**Supplementary Table 3.**
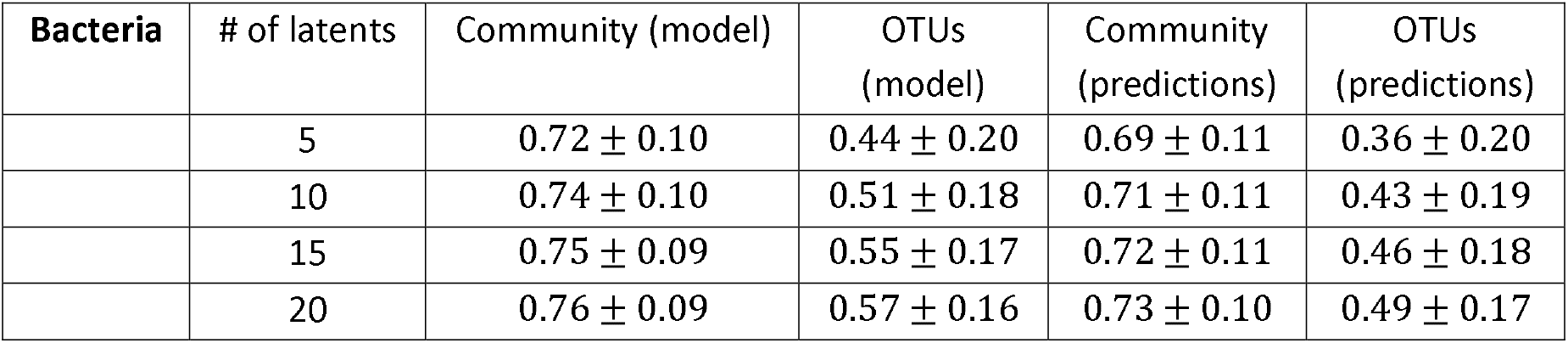
Average Spearman correlation (and standard deviation) as a function of number of latents. The parantheses “model” indicates the correlation coefficient was calculated between data and corresponding model fit. The parantheses “predictions” indicates that the correlation coefficient was calculated between testing data and model predictions. “Community” correlation coefficients measure the accuracy of reconstruction of community composition. “OTUs” correlation coefficients measure the accuracy of OTU abundance reconstruction across host samples. The numbers are shown for the bacterial community.

| Bacteria | # of latents | Community (model) | OTUs (model) | Community (predictions) | OTUs (predictions) |
| --- | --- | --- | --- | --- | --- |
| | 5 | $0.72 \pm 0.10$ | $0.44 \pm 0.20$ | $0.69 \pm 0.11$ | $0.36 \pm 0.20$ |
| | 10 | $0.74 \pm 0.10$ | $0.51 \pm 0.18$ | $0.71 \pm 0.11$ | $0.43 \pm 0.19$ |
| | 15 | $0.75 \pm 0.09$ | $0.55 \pm 0.17$ | $0.72 \pm 0.11$ | $0.46 \pm 0.18$ |
| | 20 | $0.76 \pm 0.09$ | $0.57 \pm 0.16$ | $0.73 \pm 0.10$ | $0.49 \pm 0.17$ |

**Supplementary Table 4.** Average Spearman correlation (and standard deviation) as a function of number of latents. The parantheses “model” indicates the correlation coefficient was calculated between data and corresponding model fit. The parantheses “predictions” indicates that the correlation coefficient was calculated between testing data and model predictions. “Community” correlation coefficients measure the accuracy of reconstruction of community composition. “OTUs” correlation coefficients measure the accuracy of OTU abundance reconstruction across host samples. The numbers are shown for the fungal community.

| Fungi | # of latents | Community (model) | OTUs (model) | Community (predictions) | OTUs (predictions) |
| --- | --- | --- | --- | --- | --- |
|  | 5 | 0.72 ± 0.10 | 0.45 ± 0.15 | 0.66 ± 0.18 | 0.36 ± 0.15 |
|  | 10 | 0.75 ± 0.09 | 0.53 ± 0.19 | 0.69 ± 0.16 | 0.44 ± 0.17 |
|  | 15 | 0.76 ± 0.09 | 0.55 ± 0.18 | 0.69 ± 0.14 | 0.44 ± 0.16 |
|  | 20 | 0.77 ± 0.08 | 0.57 ± 0.19 | 0.70 ± 0.16 | 0.44 ± 0.19 |

**Supplementary Table 5.** Average Spearman correlation (and standard deviation) as a function of number of latents. The parantheses “model” indicates the correlation coefficient was calculated between data and corresponding model fit. The parantheses “predictions” indicates that the correlation coefficient was calculated between testing data and model predictions. “Community” correlation coefficients measure the accuracy of reconstruction of z-scores of all metadata features in a given host sample, “metadata” correlation coefficients measure the accuracy of metadata reconstruction across host samples.The numbers are shown for the host metadata features.

| Host metadata | # of latents | Community (model) | metadata (model) | Community (predictions) | metadata (predictions) |
| --- | --- | --- | --- | --- | --- |
|  | 5 | 0.71 ± 0.11 | 0.68 ± 0.25 | 0.39 ± 0.32 | 0.40 ± 0.20 |
|  | 10 | 0.83 ± 0.08 | 0.82 ± 0.16 | 0.40 ± 0.28 | 0.41 ± 0.22 |
|  | 15 | 0.89 ± 0.05 | 0.89 ± 0.10 | 0.34 ± 0.30 | 0.37 ± 0.23 |
|  | 20 | 0.93 ± 0.04 | 0.93 ± 0.06 | 0.37 ± 0.26 | 0.38 ± 0.20 |

**Supplementary Table 6.** The L_2_-norm between the feature matrix inferred using only one farm and the projection of the same feature matrix inferred using all farms but the one farm. The brackets show the average of the same L_2_-norm calculated over 100 randomized feature matrices. Bold entries indicate statistically significant difference (evaluated using the Student’s t-test, p < 0.05)

| Farm | Archea | Bacteria | Fungi | Metadata |
| --- | --- | --- | --- | --- |
| 1 | **1.03 (3.92)** | **5.92 (7.07)** | 11.54 (7.71) | **2.49 (3.92)** |
| 2 | **1.69 (3.88)** | **4.02 (5.32)** | 9.56 (7.45) | **3.76 (4.22)** |
| 3 | **1.80 (2.98)** | **6.50 (7.34)** | 5.24 (5.20) | 2.18 (2.25) |
| 4 | **1.51 (4.81)** | **5.00 (6.05)** | **5.65 (7.00)** | **3.11 (4.58)** |
| 5 | **1.22 (3.54)** | 6.11 (5.86) | 7.43 (7.56) | **3.15 (4.06)** |

Supplementary Table 7. Table of Spearman correlation coefficients between model predictions and the corresponding metadata features. (Attached as a file)

## References

1. Gilbert, J. A. et al. Current understanding of the human microbiome. Nat. Med. 24, 392–400 (2018).

2. Borrel, G., Brugère, J.-F., Gribaldo, S., Schmitz, R. A. & Moissl-Eichinger, C. The host-associated archaeome. Not. Rev. Microbiol. 18, 622–636 (2020).

3. Huffnagle, G. B. & Noverr, M. C. The emerging world of the fungal microbiome. Trends Microbiol. 21, 334–341 (2013).

4. Chabé, M., Lokmer, A. & Ségurel, L. Gut Protozoa: Friends or Foes of the Human Gut Microbiota? Trends Porositol. 33, 925–934 (2017).

5. Liang, G. & Bushman, F. D. The human virome: assembly, composition and host interactions. Nat. Rev. Microbiol. 19, 514–527 (2021).

6. Young, V. B. The role of the microbiome in human health and disease: an introduction for clinicians. BMJ/j831 (2017) doi:10.1136/bmj.j831.

7. Tapio, I., Snelling, T. J., Strozzi, F. & Wallace, R. J. The ruminal microbiome associated with methane emissions from ruminant livestock. J. Anim. Sci. Biotechnol. 8, 7 (2017).

8. Jansson, J. K. & Baker, E. S. A multi-omic future for microbiome studies. Nat. Microbiol. 1, 16049 (2016).

9. Leibold, M. A. et al. The metacommunity concept: a framework for multi-scale community ecology: The metacommunity concept. Ecol. Lett. 7, 601–613 (2004).

10. Lynch, J. B. & Hsiao, E. Y. Microbiomes as sources of emergent host phenotypes. Science 365, 1405–1409 (2019).

11. Baxter, N. T. et al. Dynamics of Human Gut Microbiota and Short-Chain Fatty Acids in Response to Dietary Interventions with Three Fermentable Fibers. mBio 10, e02566–18 (2019).

12. Spor, A., Koren, O. & Ley, R. Unravelling the effects of the environment and host genotype on the gut microbiome. Nat. Rev. Microbiol. 9, 279–290 (2011).

13. Dubinkina, V., Fridman, Y., Pandey, P. P. & Maslov, S. Multistability and regime shifts in microbial communities explained by competition for essential nutrients. eLife 8, e49720 (2019).

14. Welsh, R. M. et al. Bacterial predation in a marine host-associated microbiome. ISME J. 10, 1540–1544 (2016).

15. ter Horst, R. et al. Host and Environmental Factors Influencing Individual Human Cytokine Responses. Cell 167, 1111–1124.e13 (2016).

16. Kleine Bardenhorst, S. et al. Data Analysis Strategies for Microbiome Studies in Human Populations—a Systematic Review of Current Practice. mSystems 6, e01154–20 (2021).

17. Shahin, M., Ji, B. & Dixit, P. D. EMBED: Essential Microbiome Dynamics, a dimensionality reduction approach for longitudinal microbiome studies. http://biorxiv.org/lookup/doi/10.1101/2021.03.18.436036 (2021) doi:10.1101/2021.03.18.436036.

18. Dixit, P. D. Thermodynamic inference of data manifolds. Phys. Rev. Res. 2, 023201 (2020).

19. The Integrative HMP (iHMP) Research Network Consortium. The Integrative Human Microbiome Project. Nature 569, 641–648 (2019).

20. Johnson, T. J. et al. A Consistent and Predictable Commercial Broiler Chicken Bacterial Microbiota in Antibiotic-Free Production Displays Strong Correlations with Performance. Appl. Environ. Microbiol. 84, e00362–18 (2018).

21. Wallace, R. J. et al. A heritable subset of the core rumen microbiome dictates dairy cow productivity and emissions. Sci. Adv. 5, eaav8391 (2019).

22. Lloyd-Price, J. et al. Multi-omics of the gut microbial ecosystem in inflammatory bowel diseases. Nature 569, 655–662 (2019).

23. Plata, G. et al. Growth promotion and antibiotic induced metabolic shifts in the chicken gut microbiome. Commun. Biol. 5, 293 (2022).

## References

1. Shahin, M., Ji, B. & Dixit, P. D. EMBED: Essential Microbiome Dynamics, a dimensionality reduction approach for longitudinal microbiome studies. http://biorxiv.org/lookup/doi/10.1101/2021.03.18.436036 (2021) doi:10.1101/2021.03.18.436036.

2. Wallace, R. J. et al. A heritable subset of the core rumen microbiome dictates dairy cow productivity and emissions. Sci. Adv. 5, eaav8391 (2019).

3. Johnson, T. J. et al. A Consistent and Predictable Commercial Broiler Chicken Bacterial Microbiota in Antibiotic-Free Production Displays Strong Correlations with Performance. Appl. Environ. Microbiol. 84, e00362–18 (2018).

4. Lloyd-Price, J. et al. Multi-omics of the gut microbial ecosystem in inflammatory bowel diseases. Nature 569, 655–662 (2019).

5. Plata, G. et al. Growth promotion and antibiotic induced metabolic shifts in the chicken gut microbiome. Commun. Biol. 5, 293 (2022).

6. Ji, B. W. et al. Quantifying spatiotemporal variability and noise in absolute microbiota abundances using replicate sampling. Nat. Methods 16, 731–736 (2019).

